# Family cluster comparisons to detect infection by *Mycobacterium leprae* in at-risk populations of six endemic regions in Colombia

**DOI:** 10.1101/2021.01.08.425863

**Authors:** Héctor Serrano-Coll, Yuliana Osorio-Leal, María Victoria Escobar-Builes, Nora Cardona-Castro

**Affiliations:** Leprosy Research Group – Colombian Institute of Tropical Medicine – CES University, Medellín, Colombia

**Keywords:** Leprosy, public health, antibodies, early detection, serologic tests, infection control

## Abstract

**Introduction:** Leprosy is a chronic infectious disease, caused by *Mycobacterium leprae*, which is endemic in some tropical countries. It is necessary to implement strategies for its detection and elimination. We propose a strategy useful could be identifying risk factors associated with a seropositive test in leprosy.

**Objective:** We aimed to quantify the infection rates and identify risk factors for M. leprae infection using the serological evaluations against NDO-LID in family clusters of leprosy patients, from regions with a high burden of leprosy in Colombia.

**Results:** We observed that belong to a low socioeconomic condition (OR 5.6 [95% IC 1.1-29]) and living in geographic area of residing such as Chocó and Atlántico (OR 2 [95% IC 1.1-3.7]) could be risk factors related to infection by *M. leprae* between the members of a family cluster.

**Conclusions:** Leprosy is a persistent disease that affects vulnerable and large family clusters, in which the detection of antibodies against NDO-LID can be a useful tool for early detection of *M. leprae* in family clusters with high risk for this infection.

## Introduction

Leprosy is a chronic granulomatous disease caused by *Mycobacterium leprae*. This mycobacterium is a Gram-positive, acid-alcohol resistant and intracellular obligate bacterium with tropism for the skin and peripheral nerves. *M. leprae* is associated with disabilities and deformities, mainly in the eyes, hands, and feet (1).

This neglected disease is a public health issue in some tropical countries despite the efforts made by the World Health Organization (WHO) for its elimination (2). In 2019, globally, 202,185 new cases of leprosy were diagnosed, and in the Americas region, 29,936 new cases of leprosy were registered (3). Furthermore, 70% of new cases of leprosy are multibacillary (MB), which represents a persistent and active transmission pathway of leprosy (4). Therefore, it is important to develop new strategies for the control of this disease.

One of these new strategies that could be key in the early detection of new cases of leprosy is the serological evaluations of NDO-LID in family clusters of leprosy patients. Given that, the serological evaluation allows to improve the follow in the household contacts (HHC) of the leprosy patients and identify whom have a higher risk to development this disease. Furthermore, this strategy will also complement the elimination goals set by WHO (5).

Therefore, the objective of this study was to quantify the infection rates and identify risk factors for M. leprae infection using the serological evaluations against NDO-LID in family clusters of leprosy patients, from regions with a high burden of leprosy in Colombia.

## Methods

### Sample study and description

This is an observational cross-sectional study, where a convenience sample of 50 family clusters was selected based on a family member diagnosed for leprosy in the last 5 years in the departments of Bolívar, Atlántico, Santander, Boyacá, Chocó and Antioquia in Colombia. The samples were obtained between 2016 and 2017. Families and patients participated voluntarily. Besides, this work was assessed and approved by the Institutional Ethics Committee for Research in Humans of CES University and Colombian Institute of Tropical Medicine.

### Sociodemographic and environmental registry

Sociodemographic and environmental characteristics were recorded through an evaluation format implemented by the Leprosy Research Group of the Colombian Institute of Tropical Medicine. This format evaluates 30 sociodemographic and environmental items related to these family clusters. In this study, we focused on the following variables: socioeconomic status, geographic location, armadillo consumption, index case classification, and number of individuals part of a family cluster.

### Measurement of antigen-specific serum antibodies (Protein A, IgM, IgG anti-NDO-LID)

Serum samples were collected from 230 members of the 50 selected family clusters for subsequent serologic evaluation. Briefly, each well of a 96-well ELISA plates (Nunc-Immuno 96-well, Polysorp plates) was coated with 1 μg/ml NDO-LID (Natural Octyl Disaccharide-Leprosy IDRI Diagnostic) antigen at room temperature and then blocked using 100 μL blocking buffer (1% bovine serum albumin [BSA] phosphate-buffered saline [PBS] +0.05%Tween 20 [PBS-T]). Plates were incubated for 1 hour with agitation at room temperature. Plates were washed (five times in PBS-T and two times in PBS), and 50 μL of serum (1:200 dilution in 0.1% BSA PBS-T) was added to each well, followed by a 1 hour of incubation with agitation of the plate at room temperature. Subsequently, 50 μL of anti-protein A; anti-IgM; anti-IgG (Rockland Immunochemicals Inc., Limerick, PA, USA) conjugated with horseradish peroxidase (HRP), diluted in 0.1% BSA PBS-T was added to each well and plates were incubated for 1 hour with agitation at room temperature. After 1 hour of incubation and washing with PBS-T, 50 μL of TMB (3,3’,5,5’-tetramethylbenzidine) substrate was added to each well and incubated for 15-minutes at room temperature. Then, the reaction was stopped by adding 25 μL 1 N sulfuric acid to each well. Optical densities (OD) were measured at 450 nm using an ELISA plate reader (Spectrophotometer Bio-Rad Xmark). Cut-off values were assessed as the average OD plus two standard deviations obtained from serum (n = 100) of healthy individuals that resided in an area not endemic for leprosy. Cut offs were assigned for anti-NDO-LID IgM (0.226 Optical densities (DO), protein A (0.127 DO), and IgG (0.183 DO).

### Statistical analysis

Data were analyzed using the Statistical Package for Social Sciences (IBM SPSS Statistics 25.0 Desktop). Univariate analysis for quantitative variables was performed through the calculation of absolute, relative frequencies and measures of central tendency. The comparison of frequencies among groups was performed using the Pearson chi-square test. Comparison of continuous value among groups was performed using the Mann-Whitney U test. The multivariate analysis was performed through a binomial logistic analysis and an approximation of the risk was made by calculating the odds ratio (OR), with its respective confidence interval (CI). P < 0.05 was considered significant.

## Results

### Sociodemographic and clinical features

A total of 50 family groups of patients with leprosy were evaluated, which were located in the departments of Bolívar and Atlántico (50%), Santander and Boyacá (30%), and Antioquia and Chocó (20%). In these family clusters, 82% were MB, and 18% were PB. When classifying them using the Ripley-Jopling scale, 58% had lepromatous leprosy, 24% had borderline leprosy, 8% had neural leprosy pure, 6% had tuberculoid leprosy and 4% had undetermined leprosy **(Table 1)**.

Regarding the socioeconomic status of these families, 58% belonged to a low socioeconomic stratum, and 42% corresponded to a medium level. Forty eight percent have a history of consuming armadillo meat. Furthermore, 72% of the clusters received Bacillus Calmette-Guerin (BCG) immunoprophylaxis. In 42% of these families, there was at least one individual with positive antibodies against *M. leprae* **(Table 1)**.

### Relation between serological data and geographic area of residence

When assessing the relationship of the serological data to the geographic area of residence, 100% of the individuals evaluated in Choco and 85.7% in the Atlantic presented at least one seropositive individual. However, we did not show statistical differences in the antibody titers between the geographic area evaluated (p> 0.005) **(Figure 1)**.

**Figure 1.**
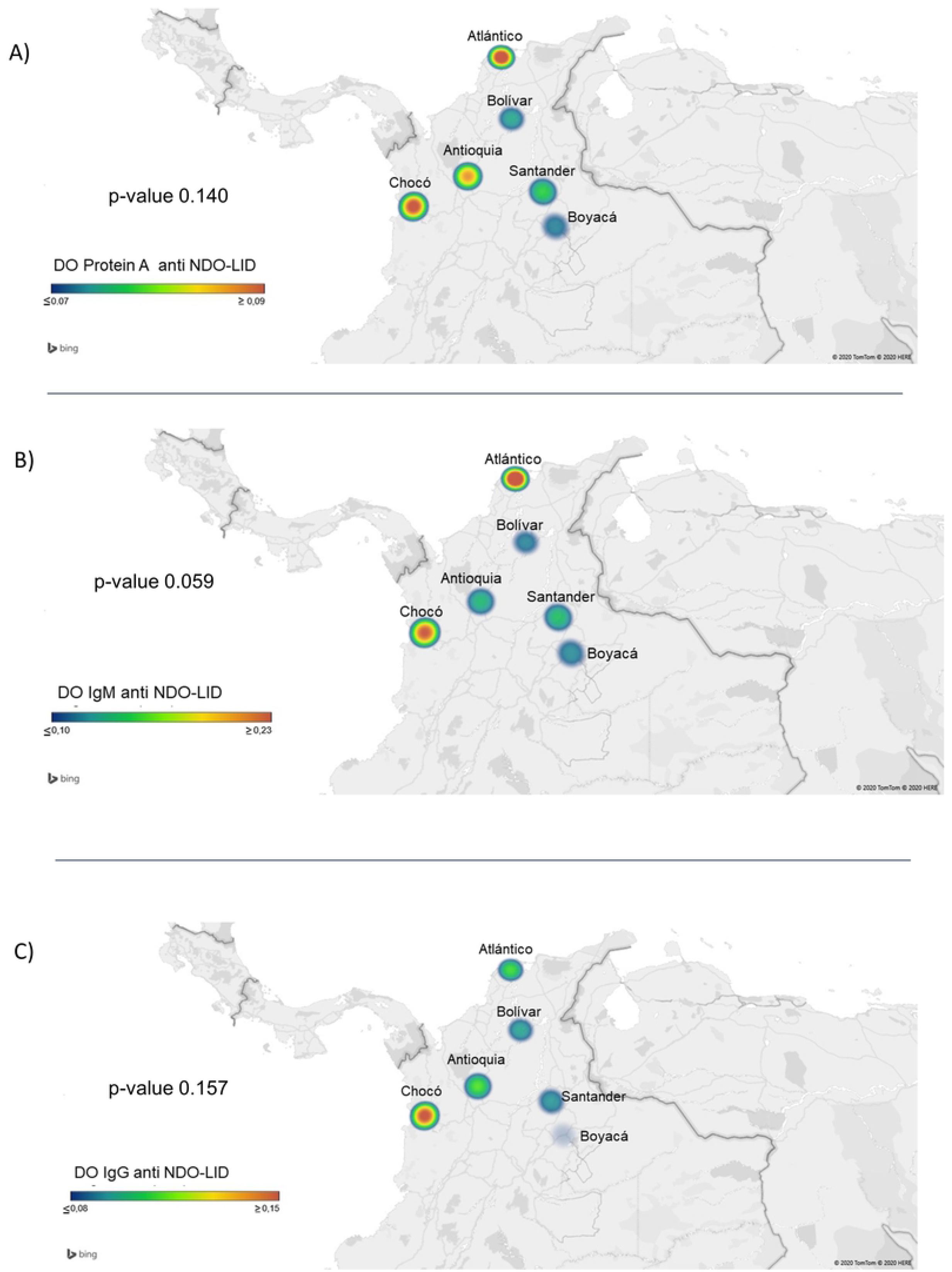

### Relation between antibody responses and socioeconomic status

When comparing the serological data with the clinical condition of these families, seropositivity, using any test, was evident in 55% of the clusters belonging to the low socioeconomic stratum. This seropositivity was statistically higher than the seropositivity of the medium socioeconomic strata (24%, p < 0.05) **(Table 3)**.

### Impact of armadillo consumption on serologic responses

Families that consumed armadillo meat (58.3%), compared to those that did not (27%), had a significantly higher risk (OR 2.1 [95% CI 1.1-4.4]) of seropositivity for any serological test. **(Table 3)**.

### Relation of serum antibody responses with the clinical form of their index case

The antibody responses in family clusters did not appear to be influenced by the clinical form of their respective index case (PB vs MB). (p> 0.05) **(Table 3)**.

### Correlation between serological responses and the number of individuals of the family cluster

When evaluating the relationship between serological responses and the number of individuals of the family cluster, a low to moderate correlation coefficient was observed for protein A (Rs Protein A = 0.28, borderline significant) and IgM (Rs IgM = 0.322, significant) **(Figure 2)**. Further, families with at least one seropositive member for any test, compared with seronegative families, had a higher median number of members per family. **(Figure 3)**.

**Figure 2.**
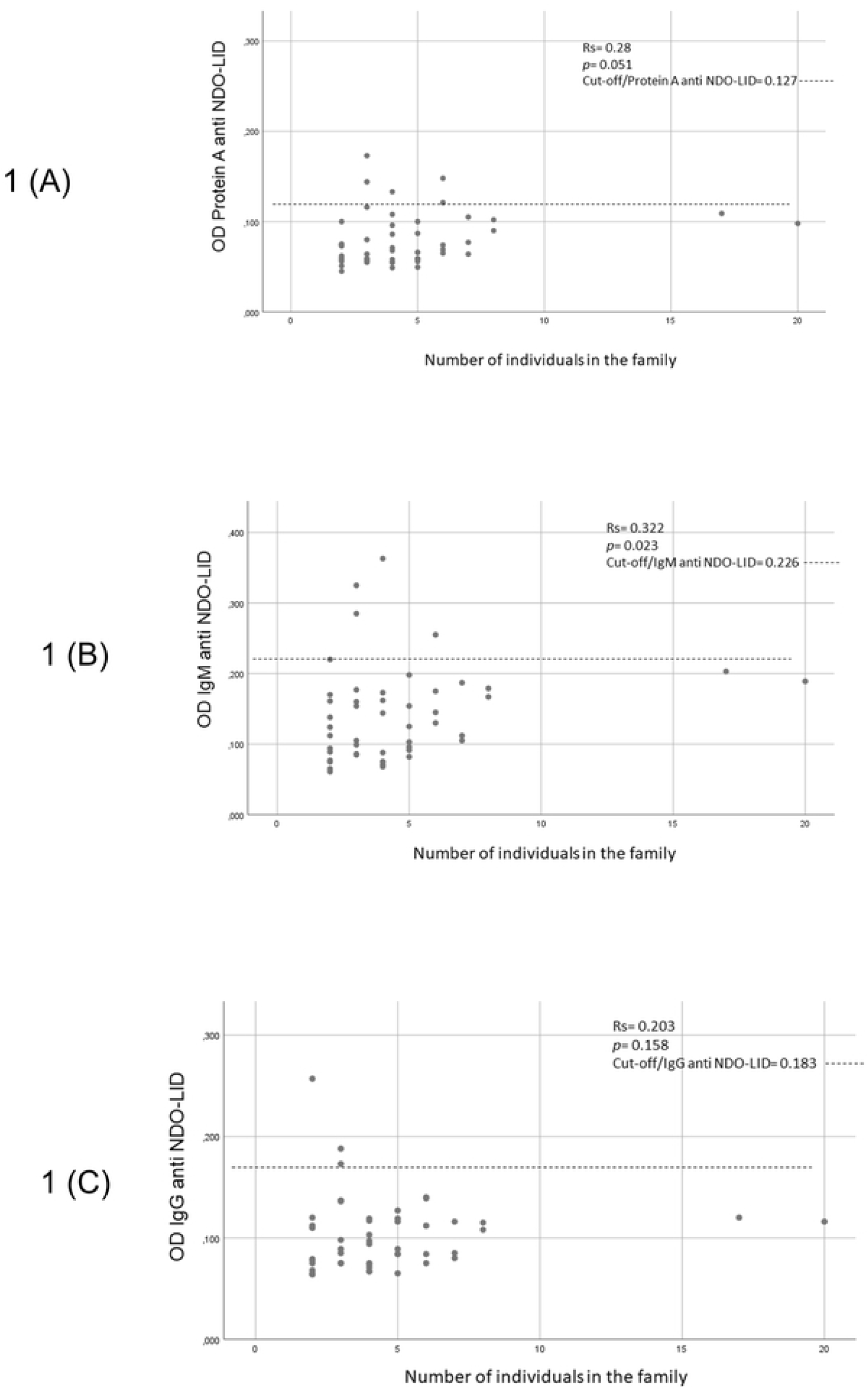

**Figure 3.**
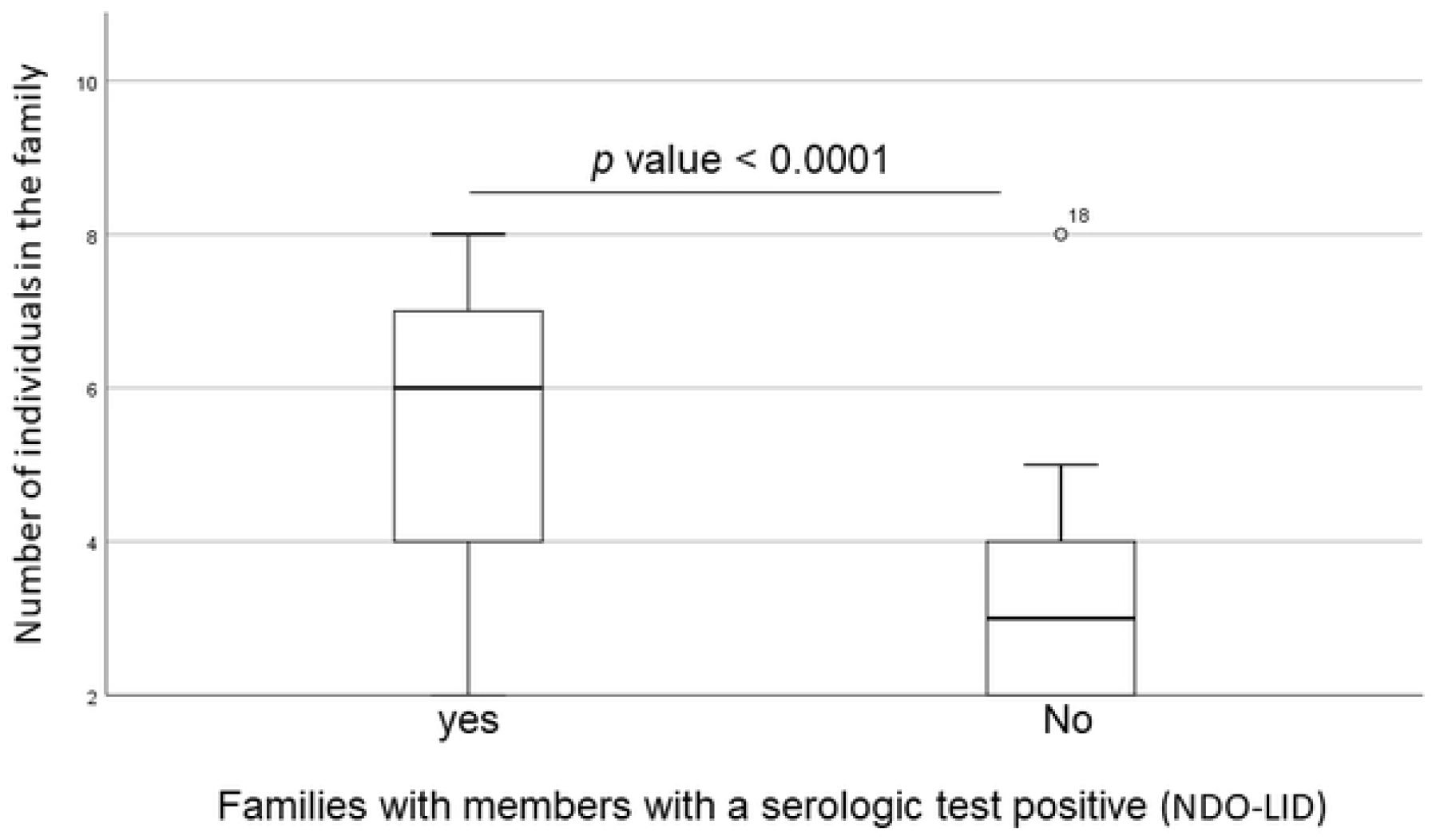

### Relation between antibody responses and social and demographic variables evaluated

A multivariate analysis of the serological data with the social and demographic variables suggested that socioeconomic condition (OR 5.6 [95% IC 1.1-29]), geographic area of residing such as Chocó and Atlántico (OR 2 [95% IC 1.1-3.7]) are risk factors to have a serological test positive (protein A) for *M. leprae* **(Table 4)**.

## Discussion

Our findings suggested a high seroconversion rate among the household contacts (HHC) of a positive *M. leprae* infected case in the Chocó and Atlántico departments. Furthermore, these results are similar to results from a study performed in Colombia, which determined the presence of positive anti-NDO-LID antibodies in healthy children and adolescents living with leprosy patients in Antioquia, Chocó, and the Caribbean regions (6).

The seropositivity observed was higher (55%) in the family clusters belonging to the low socioeconomic status. This finding suggests that *M. leprae* risk is highest for vulnerable populations, such as those in poverty (6). Populations in poverty may be at higher risk for *M. leprae* infection because many of these populations experience malnutrition and overcrowding, also risk factors for transmissibility and disease (7–10). Furthermore, this finding also agrees with other studies (11,12) including the analysis carried out in 139 municipalities of Tocantins, Brazil, where the municipalities with greater vulnerability and social inequality presented a greater number of cases of leprosy and spread of *M. leprae* (13).

This research observed that consumption of armadillo is related to higher seropositivity in the families that eaten its meat than in the family clusters that deny the consumption of this animal. This finding is similar to the results observed in a study performed in Colombia that showed higher frequencies of *M. leprae* infection in children and adolescents that ate armadillo meat (6). Furthermore, this finding is consistent with other recent research linking the consumption of armadillo meat to higher rates of leprosy cases (14). Recent result of genetic studies estimates that a third of the annual incidence of leprosy in the United States (US) may be due to contact with wild armadillos, especially in the southern region of the country (15–17). It should be noted that differentiating infection by zoonotic transmission and infection by person-to-person transmission has been complicated.

We found no relationship between PB versus MB case index status and increased likelihood for seropositivity in a family cluster. Similar findings were reported in a study carried out in Colombia in leprosy endemic areas (6). On the other hand, studies carried out in HHC of leprosy patients in Brazil reported a higher titers of anti-LID-NDO in HHC of MB patients than HHC of PB cases (9). This finding is similar to results found in other studies (18,19). However, previous research suggested that the higher titers of anti-LID-NDO is more likely related to a longer exposure time in the household (> 10 years) than to the MB link (6).

This study showed a higher seropositivity in family clusters that had a greater number of members. This finding has been previously reported in a study carried out in clusters in Indonesia, which determined that clusters with more than 7 members had a 3.1 times higher risk (95% CI: 1.3-7.3) of infection by *M. leprae* than clusters of 1-4 members (20)..

In summary, our assessment showed that the risk of *M. leprae* infection in the family clusters is increased mainly by belonging a low socioeconomic status and residing in geographic areas associated to poverty and a high burden of leprosy.

## Conclusions

This research evidenced that the set of social and demographic variables (armadillo meat intake, geographic area, low socioeconomic status, and belonging to a large family nucleus) are related to promoting seropositivity in family clusters. Furthermore, the high seropositivity in family clusters of endemic areas such as Atlántico and Chocó regions are worrying, considering that undiagnosed infected individuals may be a potential source of transmission for *M. leprae*. Therefore, in endemic areas for leprosy, control activities should be extended to the family clusters of leprosy patients, considered a high risk of infections in this population.

Finally, the serological evaluation using anti-NDO-LID could detect a significant number of patients in the early stages of this disease. Therefore, the possibility and feasibility of carrying out large-scale screening to detect antibodies against NDO-LID in family clusters of leprosy patients could be considered as a key strategy for the control and elimination of this disease in areas with a high burden of leprosy.

## Declaration of data availability

The databases generated and analyzed during this study are not publicly available but may be available upon reasonable request to the authors.

## Ethics statement

This research was carried out following international ethical standards given by the WHO and the Pan American Health Organization, supported by the Helsinki declaration, and national legislation by resolution number 008430 from 1993 of the Colombian Ministry of Health that regulates health studies. Besides, this work was assessed and approved by the Institutional Ethics Committee for Research in Humans of CES University and Colombian Institute of Tropical Medicine.

## Authors’ contributions

NCC designed the study. HSC, NCC and JCB evaluated participants in this study. JBC performed the serologic assays. HSC performed the data analysis. HSC, MVEB, NCC wrote the manuscript. All authors read and approved the manuscript.

## Acknowledgments

The authors thank to the members of the family clusters of the leprosy patients that voluntarily participated in this study and the Secretary of Health of the geographic zones evaluated. The authors also thank Dr. Luis Ernesto López and the Colombian Institute of Tropical Medicine for their collaboration in the development of this study, and Dr. Juan Leon, for editorial suggestions. We thank the Sanatorio de Contratación for the logistical support provided to this project.

## List of abbreviations

WHO: World Health Organization.
MB: Multibacillary.
PB: Paucibacillary.
NDO-LID: Natural Octyl Disaccharide-Leprosy IDRI Diagnostic.
OD: Optical densities.
OR: Odds ratio.
CI: Confidence interval.
BCG: Bacillus Calmette-Guerin.
US: United States.

